# Metagenomic Sequencing for Combined Detection of RNA and DNA Viruses in Respiratory Samples from Paediatric Patients

**DOI:** 10.1101/492835

**Authors:** Sander van Boheemen, Anneloes L. van Rijn-Klink, Nikos Pappas, Ellen C. Carbo, Ruben H.P. Vorderman, Peter J. van `t Hof, Hailiang Mei, Eric C.J. Claas, Aloys C.M. Kroes, Jutte J.C. de Vries

## Abstract

**Introduction:** Viruses are the main cause of respiratory tract infections. Metagenomic next-generation sequencing (mNGS) enables the unbiased detection of all potential pathogens in a clinical sample, including variants and even unknown pathogens. To apply mNGS in viral diagnostics, there is a need for sensitive and simultaneous detection of RNA and DNA viruses. In this study, the performance of an in-house mNGS protocol for routine diagnostics of viral respiratory infections, with single tube DNA and RNA sample-pre-treatment and potential for automated pan-pathogen detection was studied.

**Materials and Methods:** The sequencing protocol and bioinformatics analysis was designed and optimized including the optimal concentration of the spike-in internal controls equine arteritis virus (EAV) and phocine-herpes virus-1 (PhHV-1).The whole genome of PhHV-1 was sequenced and added to the NCBI database. Subsequently, the protocol was retrospectively validated using a selection of 25 respiratory samples with in total 29 positive and 346 negative PCR results, previously sent to the lab for routine diagnostics.

**Results:** The results demonstrated that our protocol using Illumina Nextseq 500 sequencing with 10 million reads showed high repeatability. The NCBI RefSeq database as opposed to the NCBI nucleotide database led to enhanced specificity of virus classification. A correlation was established between read counts and PCR cycle threshold value, demonstrating the semi-quantitative nature of viral detection by mNGS. The results as obtained by mNGS appeared condordant with PCR based diagnostics in 25 out of the 29 (86%) respiratory viruses positive by PCR and in 315 of 346 (91%) PCR-negative results. Viral pathogens only detected by mNGS, not present in the routine diagnostic workflow were influenza C, KI polyomavirus, and cytomegalovirus.

**Conclusions:** Sensitivity and analytical specificity of this mNGS protocol was comparable with PCR and higher when considering off-PCR target viral pathogens. All potential viral pathogens were detected in one single test, while it simultaneously obtained detailed information on detected viruses.

## INTRODUCTION

Respiratory tract infections pose a great burden on public health, causing extensive morbidity and mortality among patients worldwide [1–3]. The majority of acute respiratory infections is caused by viruses, such as rhinovirus (RV), influenza (INF) A and B viruses, metapneumovirus (MPV), and respiratory syncytial virus (RSV) [4]. However, in 20-62% of the patients, no pathogen is detected [4–6]. This might be the result of diagnostic failures or even infection by unknown pathogens, such as the Middle East respiratory syndrome coronavirus (MERS-CoV), that was first recognized in 2012 [7].

Rapid identification of the respiratory pathogen is critical to determine downstream decision making such as isolation measures or treatment, including cessation of antibiotic therapy. Current diagnostic amplification methods as real-time polymerase chain reaction (qPCR) are very sensitive and specific, but are aiming at particular virus species or types. Genetic diversity within the virus genome and the sheer number of potential pathogens in many clinical conditions pose limitations to predefined primer and probe based approaches, leading to false negative results [8]. These limitations, combined with the potential emergence of new or unusual pathogens highlight the need for less restricted approaches that could improve the diagnosis and subsequent outbreak management of infectious diseases.

Metagenomics relates to the study of the complete genomic content in a complex mixture of (micro)organisms [9]. Unlike bacteria, viruses do not display a common gene in all virus families, and therefore pan-virus detection relies on catch-all analytic methods. Metagenomics or untargeted next-generation sequencing (mNGS) offers a culture and nucleotide-sequence-independent method that eliminates the need to define the targets for diagnosis beforehand. Besides primary detection, mNGS immediately offers additional information, on virulence markers, epidemiology, genotyping, and evolution of pathogens [7, 10–12]. Furthermore, quantitative assessment of the presence of virus copies in the sample is enabled by the number reads [8].

While original mNGS studies typically aim at analysis of (shifts in) population diversity of abundant DNA microbes, detection of viral pathogens in patient samples requires a different technical approach because of 1) the very low abundancy of viral pathogens (<1%) in clinical samples and 2) the requisite of detection of both DNA and RNA viruses. Hence, a low limit of detection for RNA and DNA in one single assay is essential for implementation of mNGS for routine pathogen detection in clinical diagnostic laboratories. Current viral mNGS protocols are optimized for either RNA or DNA detection [11, 13–15]. Consequently, detection of both RNA and DNA viruses requires parallel work-up of both RNA and DNA pre-treatment methods. Additionally, to increase the relative concentration of viral sequences, viral particle enrichment techniques are often applied [8, 12]. These techniques are laborious and not easily automated for routine clinical diagnostic use. Moreover, during enrichment directed at viral particles, intracellular viral nucleic acids as genomes and mRNAs are being discarded. Following sequencing, the bioinformatics classification and interpretation of the results remain a major challenge. Bioinformatic classifiers are often developed for usage in either microbiome studies or classification of high abundant reads whereas extensive validation for clinical diagnostic usage in settings of very low abundancy is very limited. After bioinformatics classification, the challenge remains to discriminate between viruses that play a role in aetiology and bona fide viruses [16]. Before mNGS might be considered in routine diagnostics, there is a need for critical evaluation and validation of every step in the procedure.

In this study, we evaluated a metagenomic protocol for NGS-based pathogen detection with sample pre-treatment for DNA and RNA in a single tube. The method was validated using a selection of 25 respiratory paediatric samples with in total 29 positive and 346 negative viral PCR results. The main study objective was to define a sensitive and specific method for mNGS to be used as a broad diagnostic tool for viral respiratory diseases with the potency for automated pan-pathogen detection.

## MATERIAL AND METHODS

### Sample selection

Twenty-five stored clinical respiratory samples (−80 °C) from paediatric patients, sent to the microbiological laboratory for routine viral diagnostics in 2016, were selected from the laboratory database (GLIMS, MIPS, Belgium) at the Leiden University Medical Center (LUMC). The selection was based on respiratory virus PCR test results: 21 out of the 25 samples had one or more positive PCR result, with a variety of respiratory viruses and a wide range of quantification cycle (Cq) values. The lab-developed real-time multiplex PCR method used was an updated version of the assay previously described by Loens et al [17]. The sample types represented the routine diagnostic samples from paediatric patients sent to our laboratory: predominantly nasopharyngeal washings (n=17), sputum (n=2) and broncho-alveolar lavages (BAL, n=2). The patient selection (age range 1.2 months – 15 years) represented the paediatric population with respiratory diagnostics in our university hospital in terms of (underlying) illness.

### Sample pre-treatment

A sample pre-treatment protocol was designed with 1) potential for automation, 2) potential for pan-pathogen detection and 3) detection of intracellular viral nucleic acids. Consequently, any type of viral enrichment steps were excluded (filtration, centrifugation, nucleases, rRNA removal). Total nucleid acids (NA) were extracted directly from 200 ul of clinical material, using the MagNAPure 96 DNA and Viral NA Small Volume Kit (Roche Diagnostics, Almere, the Netherlands) with 100 μL output eluate.

### Internal controls

Clinical material was spiked with equine arteritis virus (EAV) and phocine herpesvirus 1 (PhHV1, kindly provided by prof. dr. H.G.M. Niesters, the Netherlands) as internal controls for RNA and DNA virus detection respectively. To determine the optimal concentration of the internal controls a dilution series was added to a mix of two pooled influenza A positive throat swabs (Cq 25) (PhHV1/EAV 1:100,000 1:10,000 1:1,000 1:100). Concentration was based on the number of mNGS reads (Centrifuge output as well as BLAST) in order to serve as control [19].

### Quality control

Before sequencing the DNA input concentration was measured with the Qubit (ThermoFisher Scientific, Waltham, USA), to determine whether there was sufficient DNA in the sample to obtain sequencing results. The range of DNA input for library preparation was 0.5 ng/μl for throat swabs (see reproducibility experiment) up to 300 ng/μl for bronchoalveolar lavages and sputa.

### Fragmentation

To compare the effect of different DNA fragmentation techniques, ten samples were 1) chemically fragmented using zinc (10 min.) and 2) physically fragmented using sonication with the Bioruptor^®^ pico (Diagenode, Seraing, Belgium, on/off time: 18/30s, 5 cycli) [20]. Three samples were also tested with the 3) high intensity settings of the Bioruptor^®^ pico (on/off time: 30/40s, 14 cycli).

### Library preparation

Libraries were constructed with 7μL extracted nucleic acids using the NEBNext^®^ Ultra™ Directional RNA Library Prep Kit for Illumina^®^ [21] using single, unique adaptors. This kit has been developed for transcriptome analyses. We made several adaptations to the manufacturers protocol in order to enable simultaneous detection of both DNA and RNA viruses: the following steps were omitted: Poly A mRNA capture isolation (Instruction manual NEB #E7420S/L, version 8.0, Chapter 1), rRNA depletion and DNase step (Chapter 2.1-2.4, 2.5B, 2.11A).

The size of fragments in the library was 300-700 bp. Adaptors were diluted 30 fold given the low RNA/DNA input and 21 PCR cycli were run post-adaptor ligation.

### Nucleotide Sequence Analysis

Sequencing was performed on Illumina HiSeq 4000 and NextSeq 500 sequencing systems (Illumina, San Diego, CA, USA), obtaining 10 million 150 bp paired-end reads per sample.

### Detection limit

To determine the detection limit of mNGS, serial dilutions (undiluted, 10^−1^, 10^−2^, 10^−3^, 10^−4^) of an influenza A positive sample was tested with both lab developed real-time PCR and mNGS.

### Repeatability (within run precision)

To determine the reproducibility of metagenomic sequencing an influenza A positive clinical sample (throat swab) was tested in quadruple. This sample was divided into separate aliquots, nucleic acids were extracted, library preparation and subsequent sequence analysis on the Illumina HiSeq 4000 was performed in one run.

### Bioinformatics: taxonomic classification

All FASTQ files were processed using the BIOPET Gears pipeline version 0.9.0 developed at the LUMC [22]. This pipeline performs FASTQ pre-processing (including quality control, quality trimming and adapter clipping) and taxonomic classification of sequencing reads. In this project, FastQC version 0.11.2 [23] was used for checking the quality of the raw reads. Low quality read trimming was done using Sickle [24] version 1.33 with default settings. Adapter clipping was performed using Cutadapt [25] version 1.10 with default settings. Taxonomic classification of reads was performed with Centrifuge [26] version 1.0.1-beta. The pre-built NT index, which contains all sequences from NCBI’s nucleotide database, provided by the Centrifuge developers was used (ftp://ftp.ccb.jhu.edu/pub/infphilo/centrifuge/data/nt.tar.gz) as the reference database.

In addition, a customized reference centrifuge index with sequence information obtained from the NCBI’s RefSeq [27] (accessed November 2017) database was built. RefSeq genomic sequences for the domains of bacteria, viruses, archaea, fungi, protozoa, as well as the human reference, along with the taxonomy identifiers, were downloaded with the Centrifuge-download utility and were used as input for centrifuge-build.

Centrifuge settings were evaluated to increase the sensitivity and specificity. The default setting, with which a read can be assigned to up to five different taxonomic categories, was compared to one unique assignment per read [26] where a read is assigned to a single taxonomic category, corresponding to the lowest common ancestor of all matching species.

Kraken-style reports with taxonomical information were produced by the Centrifuge-kreport utility for all (default) options. Both unique and non-unique assignments can be reported, and these settings were compared. The resulting tree-like structured, Kraken-style reports were visualized with Krona [28] version 2.0.

In silico simulated EAV reads were analysed in different databases (NCBI nucleotide vs RefSeq), classification algorithms (max 5 labels per sequence, vs unique (common ancestor)) and reporting (non-unique vs unique) to determine the most sensitive an specific bioinformatic analyses using Centrifuge.

To determine the amount of reads needed, results of 1 and 10 million reads were compared. 1 million reads were randomly selected of the 10 million reads of one FASTQ file and analysed.

### Bioinformatics: assembly of PhHV1 sequences

Assembly of PhHV1 was done using the biowdl virus-assembly pipeline 0.1 [29]. The QC part of the biowdl pipeline determines which adapters need to be clipped by using FastQC version 0.11.7 [23] and cutadapt version 1.16 [25], with minimum length setting “1”. The resulting reads were downsampled within bowdl to 250 000 reads using seqtk 1.2 [30] after which SPADES version 3.11.1 [31] was run to get the first proposed genome contigs.

To retrieve longer assembly contigs a reiterative assembly approach was used by processing the proposed contigs by the biowdl reAssembly pipeline 0.1. This preassembly pipeline aligns reads to contigs of a previous assembly, then selects the aligned reads, downsamples them and runs a new assembly using SPADES. Subtools used for this consisted of BWA 0.7.17 [32] for indexing and mapping, SAMtools 1.6 [33] for creating bam files, SAMtools view (version 1.7) for filtering out unmapped reads using the setting “-G 12”, Picard SamToFastq (version 2.18.4) and seqtk for creating fastq files with 250 000 reads. The contigs from the reAssembly pipeline were then processed for a second using SPADES, with setting the ‘cov-cutoff to 5. The resulting contigs were then processed with the reAssembly pipeline for the third and last time setting the ‘cov-cutoff in SPADES to 20.

The contigs from the last reAssembly step were then run against the blast NT database using blastn 2.7.1 [19] Out of 23 contigs only 5 contigs, that showed the lowest % in identity matches with any other possible non herpes virus species, were selected. The final 5 contigs contained sequence lengths of 97893, 8170 3710, 3294 and1279 nucleotides, the average coverage was 206, 131, 211, 285 and 154, respectively. These five contigs were published as partial genome under accession number (NCBI accession number: MH509440)

### Retrospective validation

Sensitivity and specificity of the metagenomic NGS procedure was compared with the lab developed multiplex qPCR [17] using a selection of 21 samples positive for at least one respiratory PCR target and 4 negative samples.. The routine multiplex PCR panel consisted of 15 respiratory target pathogens: influenza virus A/ B, respiratory syncytial virus (RSV), metapneumovirus (MPV), adenovirus (ADV), human bocavirus (HBoV), parainfluenza viruses (PIV) 1/ 2/ 3/ 4, rhinovirus (RV), and the coronaviruses HKU1, NL63, 227E and OC43. Thus, in total 375 PCR results were available (15 targets × 25 samples) of which 29 PCR positive and 346 PCR negative for comparison with mNGS. Validation samples were tested with mNGS, using the total NA extraction protocol, the adapted NEBNext library preparation protocol, and sequencing 10 million reads on an Illumina NextSeq 500. Bioinformatics analyses was performed using Centrifuge with the RefSeq database and unique assignment of sequence reads.

### Ethical approval of patient studies

The study design was approved by the medical ethics review committee of the Leiden University Medical Center.

## RESULTS

### Internal controls

Serial dilutions of EAV and PhHV1 were added to an influenza A PCR positive sample. Serial dilution 1:10,000 detected EAV with a substantial read count in the presence of a viral infection and without a significant decline in target virus family reads (Table 1). Based on these results we determined the concentration of internal controls for further experiments.

**Table 1.** Internal controls EAV/PhHV-1: serial dilutions against a clinical sample background and within-run precision (INFA)

| **Sample nr** | **INFA Cq** | **EAV Cq** | **PhHV-1 Cq** | **INFA reads centrifuge** | **INFA reads BWA** | **EAV reads Centrifuge** | **EAV reads BWA** | **PhHV-1 reads BWA** |
| --- | --- | --- | --- | --- | --- | --- | --- | --- |
| **1** | 24.79 | 30.85 | 32.55 | 8843 (3.9 log) | 9805 | 9 | 72 | 3302 |
| **2** | 24.76 | 28.45 | 30.33 | 11011 (4.0 log) | 12570 | 49 | 417 | 3773 |
| **3** | 24.67 | 24.91 | 26.83 | 8525 (3.9 log) | 9260 | 520 | 5130 | 6664 |
| **4** | 24.52 | 21.59 | 23.52 | 10010 (4.0 log) | 11384 | 5634 | 54366 | 19984 |

Abbreviations: nr: number, Cq: quantification cycle value, INFA: influenza A, EAV: equine arteritis virus, PhHV-1 phocine herpesvirus 1.
PhHV-1 reads were based on BWA alignment, since the PhHV-1 genome was lacking in the NCBI database

The EAV Cq value of the dilutions correlated with the number of EAV reads from both BLAST alignment and the Centrifuge analysis (Figure 1).

**Figure 1.**
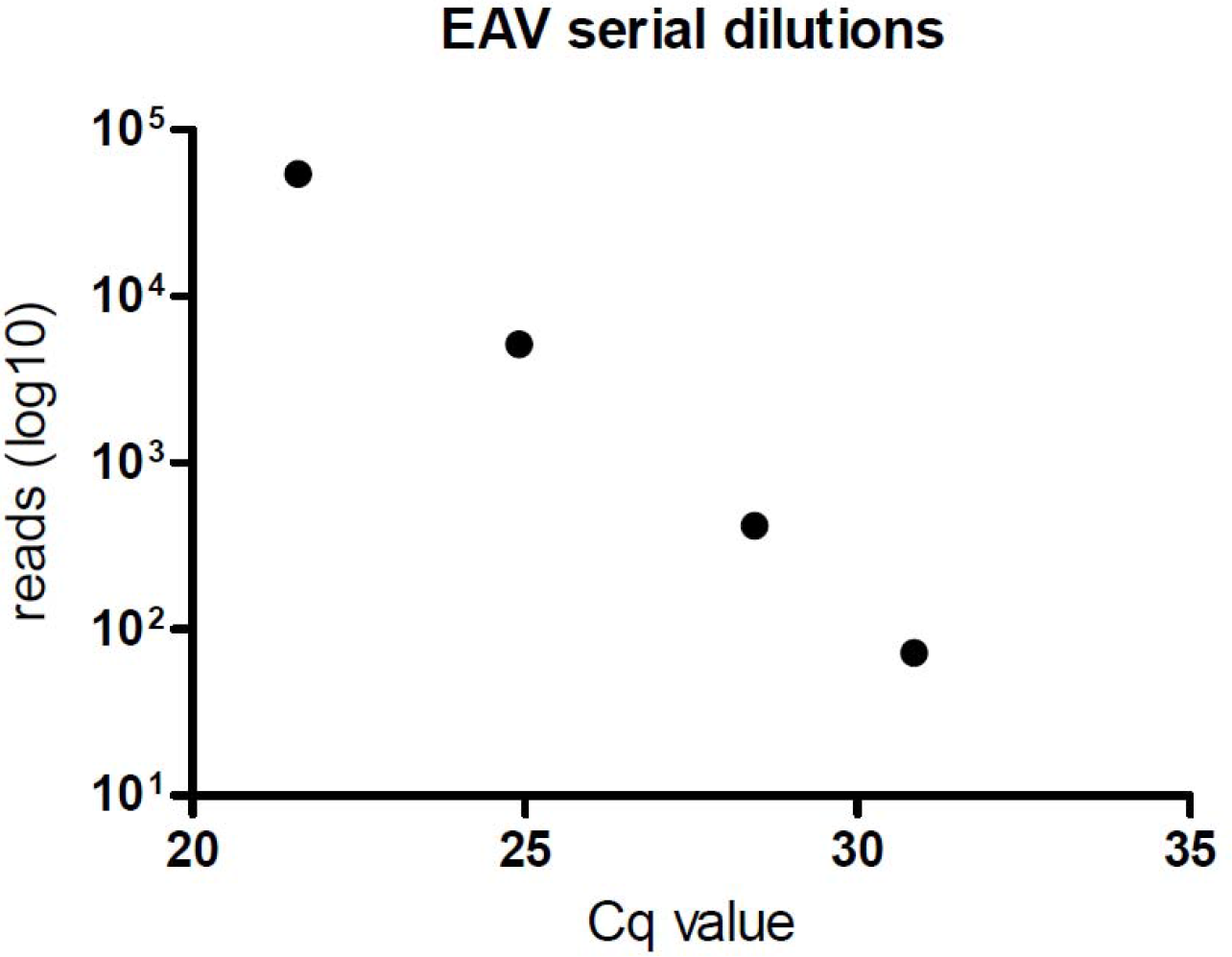
Correlation of Cq value and the number of EAV reads (serial dilutions).

Since the NCBI database was lacking a complete PhHV1 genome sequence, PhHV1 was sequenced and based on the gained sequence reads the genome was build using SPAdes [31]. The proposed almost complete genome of PhHV1 was added to the NCBI genbank database (submission ID 2124975, GenBank MH509440, release date 4 Dec 2018) and used for BLAST alignment.

### Fragmentation

The comparison of fragmentation methods for a selection of the samples with relevant target reads, is shown in Figure 2. Root reads were comparable among the three protocols. The protocol with fragmentation with Zinc had higher yield in target virus reads for all RNA viruses tested and adenovirus.

**Figure 2.**
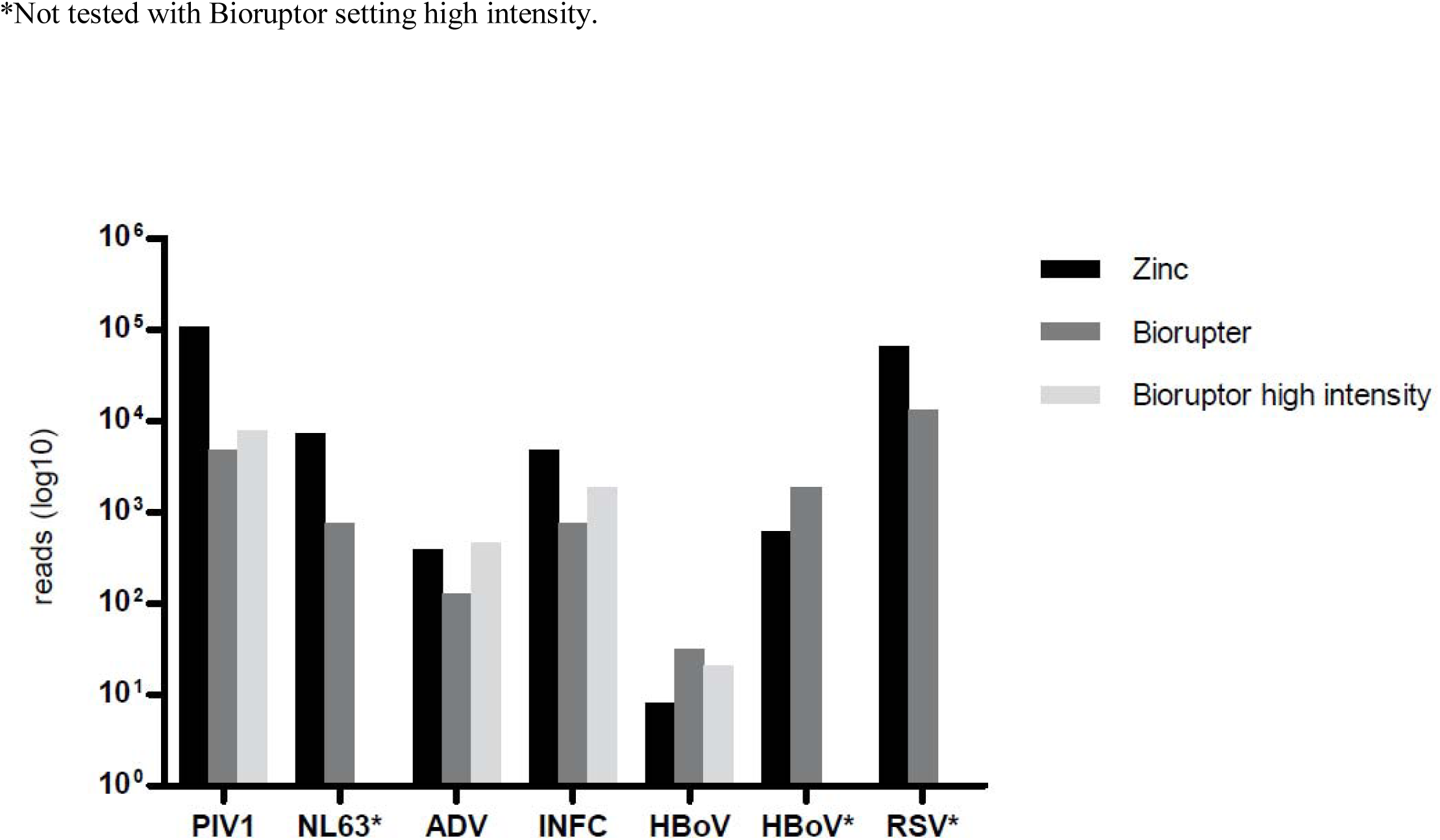
Comparison of fragmentation methods on target reads (species level, log scale).

### Index hopping

Sequence analysis by Illumina HiSeq 4000 with single, unique indexes resulted in HRV-C sequences (22-159 reads), in all samples run on the same lane, in contrast to samples run on another lane. Comparison of HRV-C sequences between these samples resulted in an exact match. Retesting of these samples with Illumina Nextseq 500 resulted in disappearance of HRV reads in the samples, with the exception of a few HRV PCR positive samples (Figure 3). Combined, this was highly suggestive for index hopping at the Illumina HiSeq 4000 [34].

**Figure 3.**
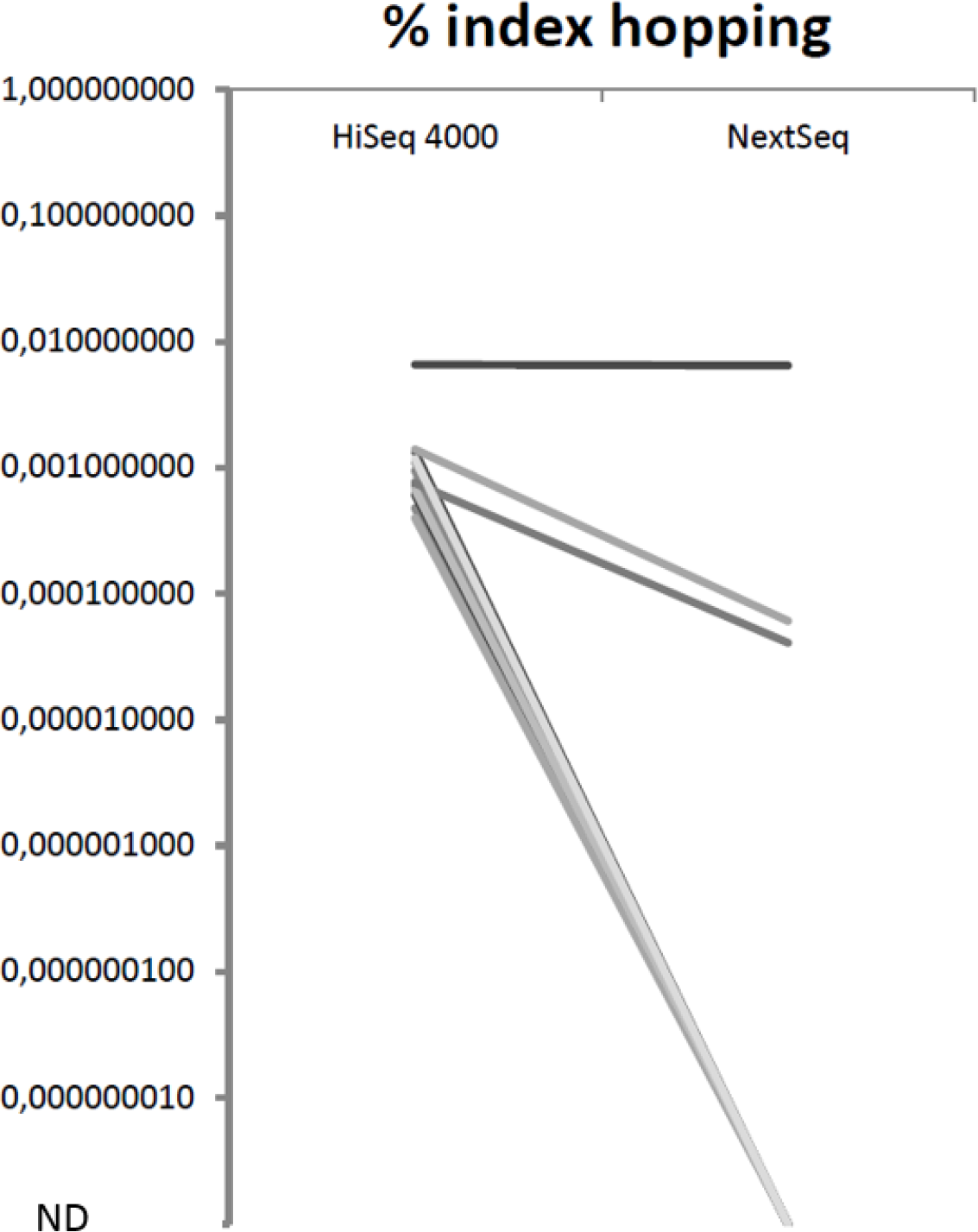
Decline in index hopping (percentages Rhinovirus C of root reads) with Illumina NextSeq 500 as compared to HiSeq 4000. Each line represents one sample.

### Detection limit

The detection limit, deduced from serial dilutions of EAV (Figure 1) and influenza A (Figure 4) was comparable with a real time PCR Cq value of approximately 35.

**Figure 4.**
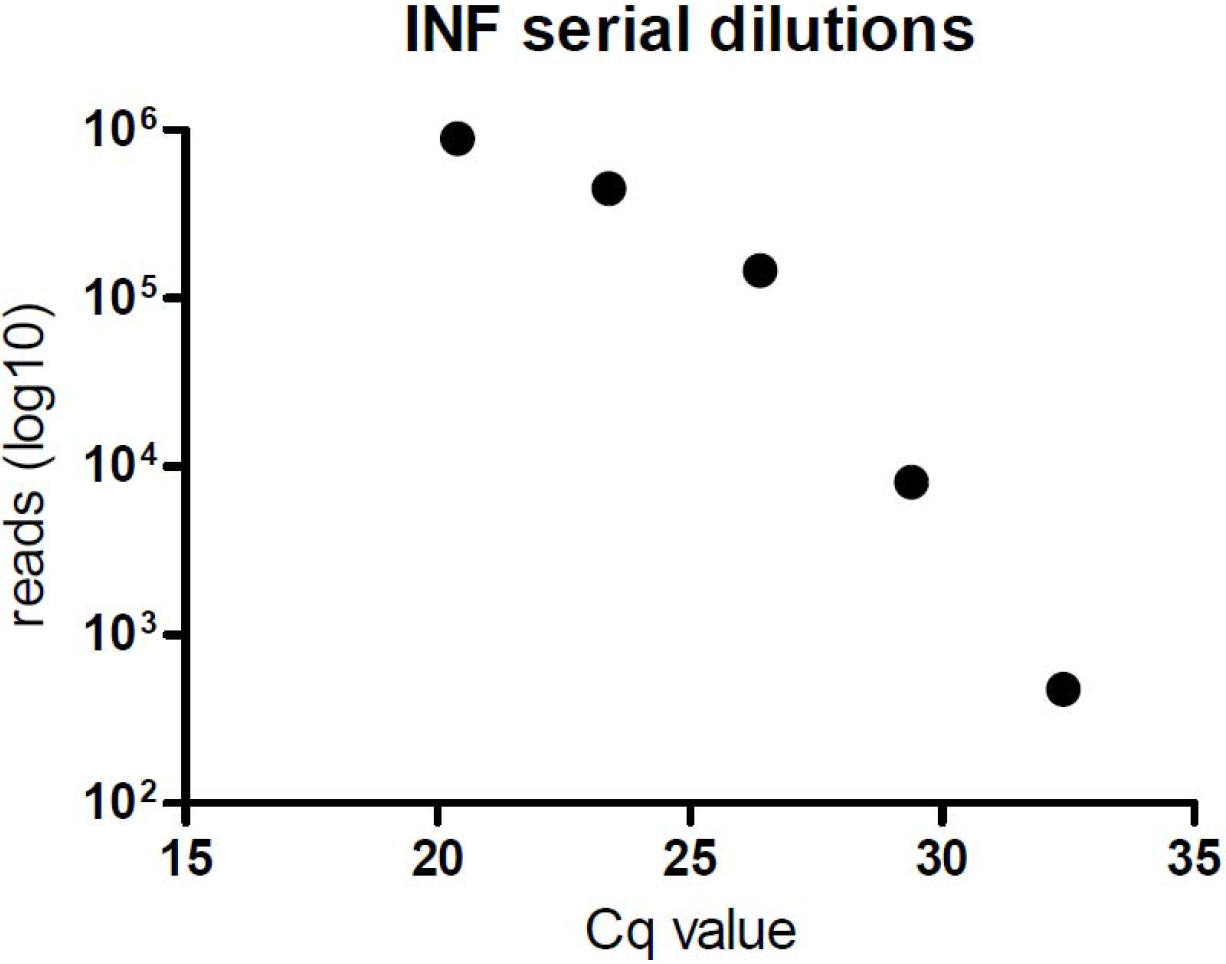
Serial dilutions of an influenza A positive clinical sample.

### Repeatability: within run precision

The mNGS results of an influenza A positive sample tested in quadruple could be reproduced with only minor differences (table 1): coefficient of variation of 1.2%: 0.05 log SD/ 4.0 log average.

### Bioinformatics: taxonomic classification

The Centrifuge default settings, with the NCBI nucleotide database and maximum 5 labels per sequence, resulted in various spurious classifications (Figure 5), for example Lassa virus (Figure 6), evidently highly unlikely to be present in patient samples from the Netherlands with respiratory complaints. The specificity could be increased by using the NCBI RefSeq database instead of the nucleotide database. The classification was further improved by changing the Centrifuge tool settings to limit the assignment of homologous reads to the lowest common ancestor (maximum 1 label per sequence). Classification with maximum 5 labels per read resulted in two different outcomes using the report with all mappings and the report with unique mappings, with the latter missing the reads assigned to multiple organisms.

**Figure 5.**
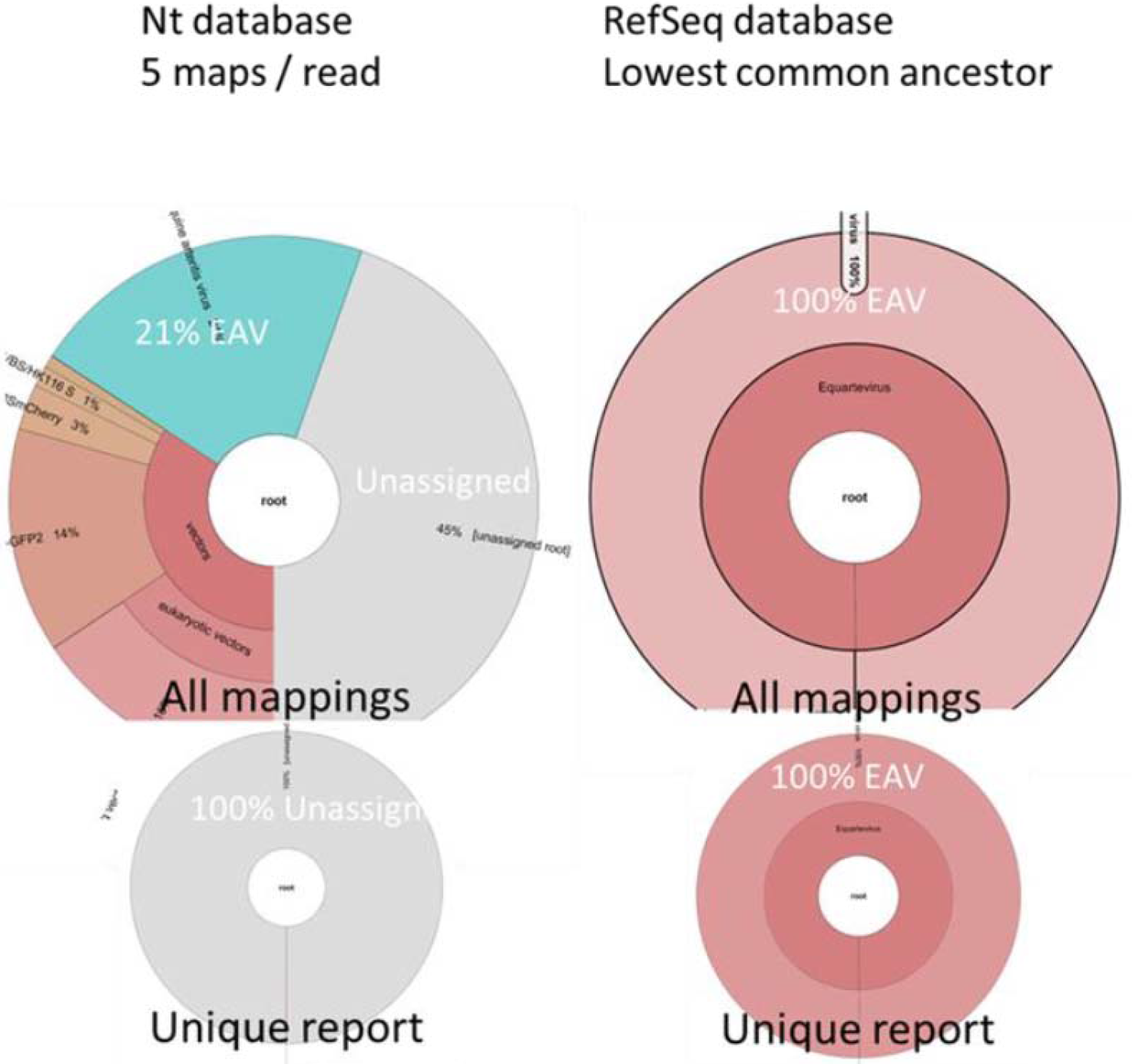
Analysis of in silico simulated EAV reads with the different bioinformatic settings of the Centrifuge pipeline.

**Figure 6.**
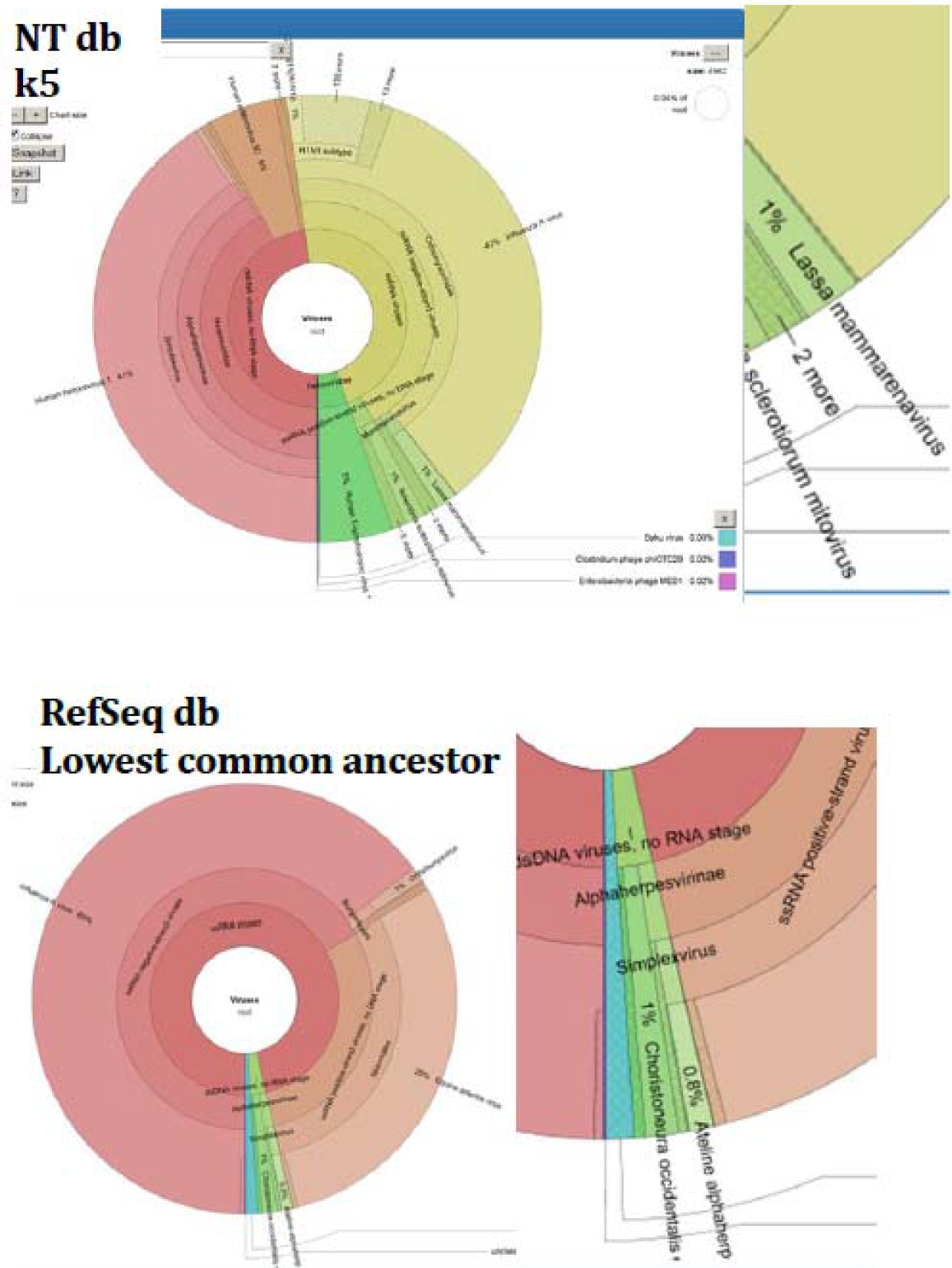
Spurious Lassa virus reads with the NCBI Nucleotide (NT) database versus the RefSeq database. k5; up to 5 labels per sequence.

Comparison of classification using these different settings shows the highest sensitivity and specificity using the RefSeq database with one label (lowest common ancestor) assignment, both with in silico prepared datasets containing solely EAV sequence fragments (Figure 5) and with clinical datasets (with highly abundant background) (Figure 6).

To determine the effect of the total number of sequencing reads obtained per sample on sensitivity, one million and 10 million reads were compared by means of in silico analysis (Table 2).

**Table 2.** Comparison of analysis of 1 million vs 10 million reads.

| virus | virusfamily | 10 million reads |  |  |  |  | 1 million reads |  |  |  |
| --- | --- | --- | --- | --- | --- | --- | --- | --- | --- | --- |
|  |  | Cq<br>value | Root<br>reads | virus<br>family<br>reads | %<br>root | %<br>viral | Root<br>reads | virus<br>family<br>reads | %<br>root | % viral |
| **RV** | Picornaviridae | 37.7 | 8203894 | 5290 | 0.06 | 84.37 | 822218 | 527 | 0.07 | 86.11 |
| **PIV4** | Paramyxoviridae | 24.9 | 10886798 | 3965 | 0.04 | 41.90 | 1088067 | 369 | 0.08 | 40.73 |
| **CMV** | Herpesviridae | 34.5 | 15889428 | 806 | 00.01 | 10.88 | 1588922 | 82 | 0.04 | 11.87 |
| **ADV** | Adenoviridae | 30.2 | 11146488 | 0 | 0 | 0 | 1115135 | 0 | 0.03 | 0 |
| **RSV** | Paramyxoviridae | 27.3 | 10191995 | 2287 | 0.02 | 53.29 | 1019415 | 253 | 0.04 | 59.25 |
| **INFB** | Orthomyxoviridae | 30 | 8535672 | 804 | 0.01 | 48.67 | 853149 | 75 | 0.02 | 46.58 |
| **NL63** | Coronaviridae | 36.2 | 10386928 | 0 | 0 | 0 | 1038469 | 0 | 0.02 | 0 |
| **INFA** | Orthomyxoviridae | 27.5 | 10981601 | 12539 | 0.11 | 70.28 | 1097872 | 1276 | 0.17 | 69.84 |
| **MPV** | Paramyxoviridae | 34.1 | 12972626 | 3 | 0 | 0.10 | 1297151 | 0 | 0.02 | 0 |
| **HBOV** | Parvoviridae | 32.2 | 11819805 | 0 | 0 | 0 | 1181738 | 0 | 0 | 0 |
| **RV** | Picornaviridae | 23.1 | 11819805 | 50034 | 0.42 | 84.27 | 1183738 | 4912 | 0.49 | 84.25 |

Abbreviations: Cq: quantification cycle value, % root: percentage of total root reads, % viral: percentage of all viral reads , RV: rhinovirus,
PIV4: parainfluenza 4, CMV: cytomegalovirus, ADV: adenovirus, RSV: respiratory syncytial virus, INF: influenza, NL63: coronavirus NL63,
MPV: metapneumovirus, hBoV: human bocavirus

### Retrospective validation

#### Clinical sensitivity based on PCR target pathogens

The sample collection consisted of 21 clinical specimens positive for at least one of the following PCR target viruses: rhinovirus, influenza A&B, parainfluenza 1 &4 (PIV), metapneumovirus, respiratory syncytial virus, coronaviruses NL63 and HKU1 (CoV), human bocavirus (hBoV), and adenovirus (ADV). Fourteen samples were positive for one virus, six samples for two and one sample for three viruses with the lab-developed respiratory multiplex qPCR. Cq values ranged from Cq 17 to Cq 35, with a median of 23.

With mNGS 25 of the 29 viruses demonstrated in routine diagnostics were detected (Table 3), resulting in a sensitivity of 86% for PCR targets. If a cut-off of 15 reads was handled, sensitivity declined to 67% (Table 4).

mNGS target read count showed a correlation with the Cq values of the qPCR (Figure 7).

**Table 3.**
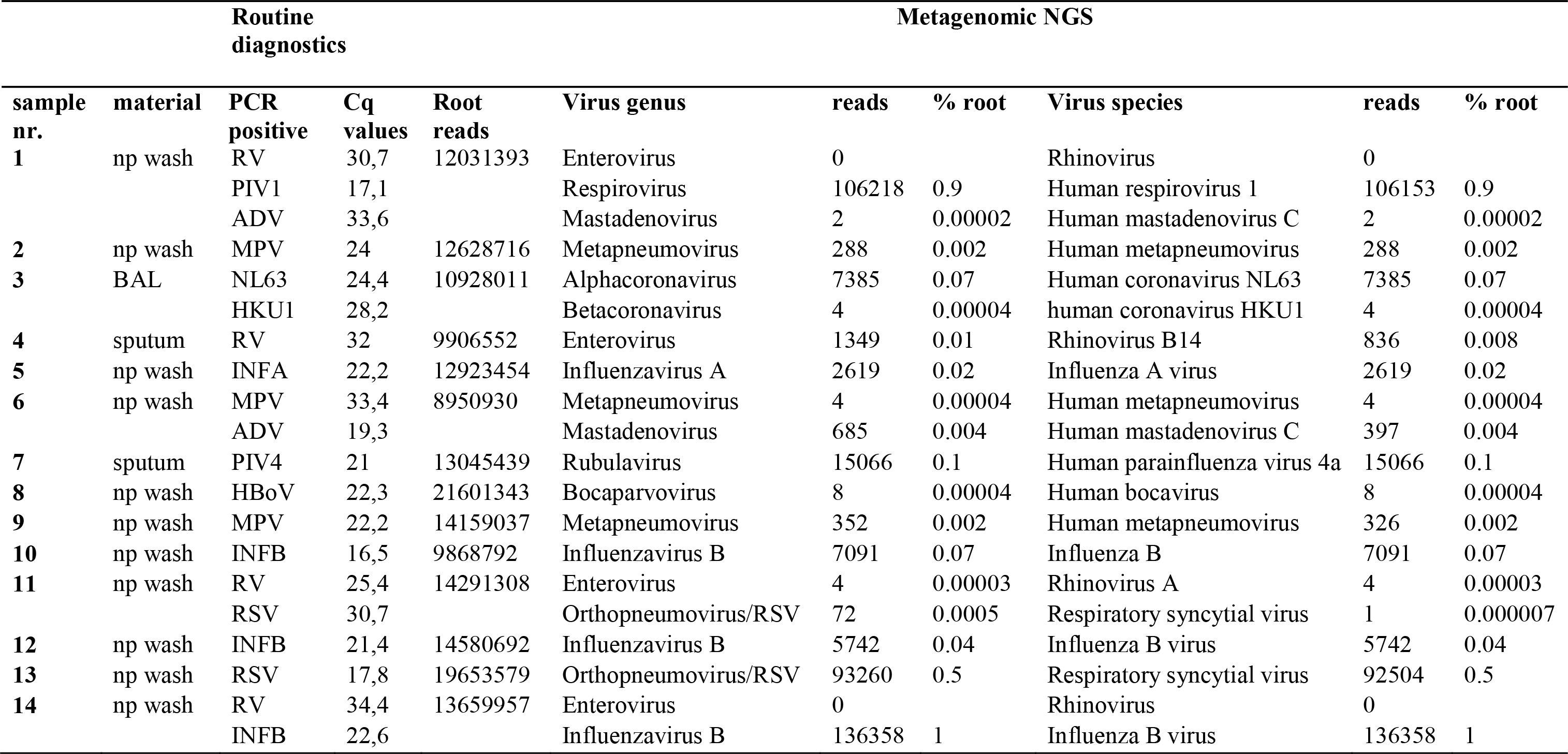
Detection of qPCR viruses positive respiratory samples with mNGS

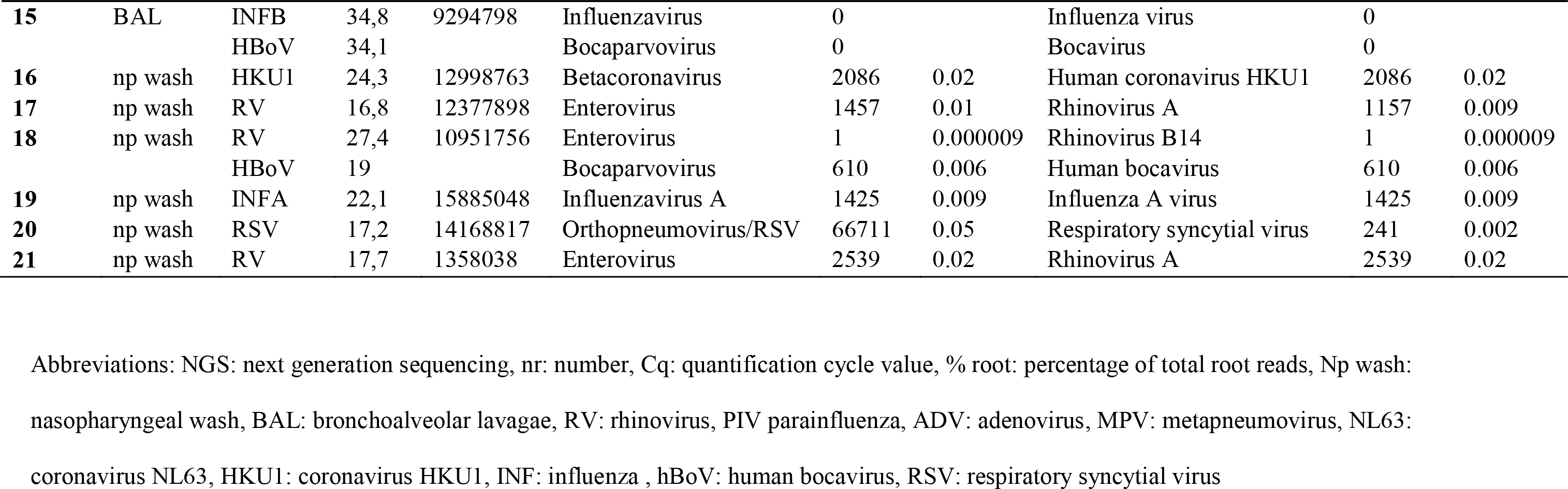

**Table 4.** Sensitivity and specificity of the mNGS protocol tested, based on PCR target virus, and different cut-off levels for defining a positive result.

|  | **All**<br>**reads** | **≥15**<br>**reads** | **≥50**<br>**reads** |
| --- | --- | --- | --- |
| **Sensitivity** | 86 | 67 | 67 |
| **Specificity** | 91 | 99 | 100 |

**Figure 7.**
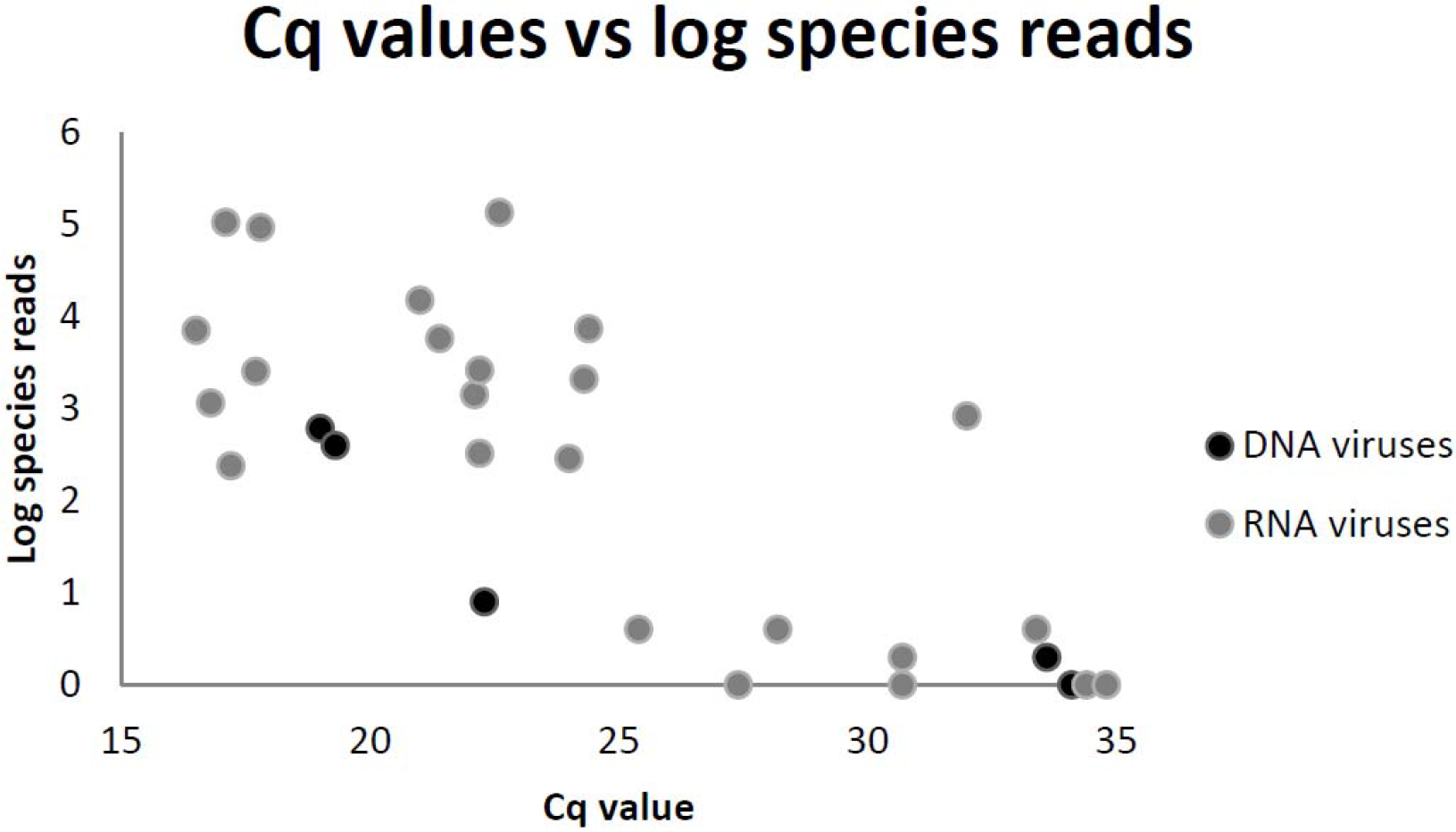
Semi-quantification of the mNGS assay for target virus detection in clinical samples with qPCR confirmed human respiratory viruses.

#### Detection of additional viral pathogens by mNGS: off-PCR target viruses

Next to the viral pathogens tested by PCR, mNGS also detected other pathogenic viruses, indicating additional viral sequences uncovered by mNGS but not included in the routine diagnostics, with influenza C virus being the most prominent. A high amount,4800 reads, of influenza virus C reads (C/Ann Arbor/1/50) (88% of all viral reads and 0.02 of the root reads) was found in one sample. Other potential respiratory pathogens detected by mNGS and not included in PCR analysis were KI polyomavirus (2 samples: 159 and 30 reads respectively), cytomegalovirus (human betaherpesvirus 5) (1704 and 132 reads). All of these viruses are not included routinely in the diagnostic multiplex qPCRs.

#### Internal controls

The spiked-in internal controls were detected by mNGS in all samples. EAV sequence reads ranged from 18-34660 (median 538) and herpesviridae reads as indicator of PhHV1 ranged from 1-1707 (median 23).

#### Analytical specificity based on PCR target viruses

In total 25 paediatric respiratory samples were available for analysis of analytical specificity of mNGS: 4 samples were negative for all 15 viral pathogens in the multiplex PCR panel (influenza A/B, RSV, HMPV, ADV, HBoV, PIV1/2/3/4, RV, HKU1, NL63, 227E, OC43) and 21 samples were negative for 12-14 of these PCR target pathogens.

Out of in total 346 negative target PCR results of these 25 samples, 315 results corresponded with the finding of 0 target specific reads by mNGS. If a cut-off of 15 reads was used 343 of the 346 negative PCR targets were negative with mNGS. The 3 samples positive by mNGS and negative by PCR were human parainfluenzavirus 1 (27 reads), 3 (31 reads), and 4 (27 reads). Though no conclusive proof for neither true positive or false mNGS results could be found, specificity of mNGS was 91% (315/346) when encountering all reads and ≥99% (343/346) with a 15 reads cut-off (Table 6).

#### Drug resistance data

Using the sequence data we identified several mutations on the genome of two clinical samples tested positive for influenza A virus. None of these mutations conferred resistance to either oseltamivir or zanamivir. For the positions where resistance associated mutations occur, amino acids I117, E119, D198, I222, H274, R292, N294 and I314 showed susceptibility to oseltamivir and V116, R118, E119, Q136, D151, R152, R224, E276, R292 and R371 revealed susceptibility to zanamivir [35, 36].

## DISCUSSION

Metagenomic sequencing has not yet been implemented as routine diagnostic tool in clinical diagnostics of viral infections. Such application would require the careful definition and validation of several parameters to enable the accurate assessment of a clinical sample with regard to the presence or absence of a pathogen, in order to fulfil current accreditation guidelines. For this purpose, this study has initiated the optimization of several steps throughout the pre- and post-sequencing workflow, which are considered essential for sensitive and specific mNGS based virus detection. Many virus discovery or virus diagnostic protocols have focussed on the enrichment of viral particles [37] with the intention to increase the relative amount of virus reads. However, these methods are laborious and intrinsically exclude viral nucleic acid located in host cells. The current protocol enabled high throughput sample pre-treatment by means of automated NA extraction and without depletion of bacterial nor human genome, with potential for pan-pathogen detection. Several adaptations in the bioinformatic script resulted in more accurate classification output reporting.

The addition of an internal control to a PCR reaction is widely used as an quality control of a qPCR [38]. While the addition of internal controls in mNGS is not yet an accepted standard procedure, we employed EAV and PhHV1 as an RNA and DNA controls, respectively, as for diagnostic application such precautions are required. The amount of internal control reads and target virus reads have been reported to be dependent of the amount of background reads (negative correlation) [39]. In our protocol, the internal controls were used as qualitative controls but may be used as indicator of the amount of background. Since the NCBI database was lacking a complete PhHV1 genome, the Centrifuge index building and classification was limited to classification on a higher taxonomic rank. In order to achieve classification of PhHV1 at species level, the whole genome of PhHV1 was sequenced, and based on the gained sequence reads the genome was build [31]. The proposed almost complete genome of PhHV1 was added to the NCBI GenBank database.

Sensitivity of the mNGS procotol was 86% (25/29) based on PCR target viruses. Four, all RNA viruses, that were not recovered by mNGS had high Cq values, over 31, i.e. a relatively low viral load. This may be a drawback of the retrospective nature of this clinical evaluation as RNA viruses may be degraded due to storage and freeze-thaw steps, resulting in lower sensitivity of mNGS. A correlation was found between read counts and PCR Cq value, demonstrating the quantitative nature of viral detection by mNGS. Discrepancies between the Cq values and the number of mNGS reads may be explained by 1) unrepresentative Cq values, e.g. by primer mismatch for highly divergent viruses like rhino/enteroviruses and 2) differences in sensitivity of mNGS for several groups of viruses, as has been reported by others [42]. Additionally, several viral pathogens were detected that were not targeted by the routine PCR assays, including influenza C virus, which is typical of the unbiased nature of the method. In addition, though not within the scope of this study, bacterial pathogens, including Bordetella pertussis (qPCR positive), were also detected. In the current study only viruses were targeted since these could be well compared to qPCR results, bacterial targets remain to be studied in clinical sample types more suitable for bacterial detection than nasopharyngeal washings. The analytical specificity of mNGS appeared to be high, especially with a cut-off of 15 reads. However, the clinical specificity, the relevance of the lower read numbers, still needs further investigation in clinical studies.

Sequencing using Illumina HiSeq 4000 chemistry resulted in frequent false positive rhinovirus reads in numerous samples. This problem could be attributed to ‘index hopping’ (index miss-assignment) as earlier described [34]. Although the percentage of reads which contributed to the index hopping was very low, this is critical for clinical viral diagnostics, as this is aimed specifically at low abundancy targets [34, 40].

Bioinformatics classification of metagenomic sequence data with the pipeline Centrifuge required identification of the optimal parameters in order to minimize miss- and unclassified reads. Default settings of this pipeline resulted in higher rates of both false positive and false negative results. The nucleotide database includes a wide variety of unannotated viral sequences, such as partial sequences and (chimeric) constructs, in contrast to the curated and well-annotated sequences in the NCBI Refseq database, which resulted in a higher specificity. In addition to the database, settings for the assignment algorithm were adapted as well. The assignment settings were adjusted to unique assignment in the case of homology to the lowest common ancestor. This modification resulted in higher sensitivity and specificity than the default settings, however the ability to further subtyping diminished. This is likely to be attributed to the limited representation/availability of strain types within the RefSeq database. In consequence, this leads to a more accurate estimation of the common ancestor for particular viruses, but limited typing results in case of highly variable ones. To obtain optimal typing results, additional annotated sequences may be added or a new database should be build, with a high variety of well-defined and frequently updated virus strain types.

To conclude, this study contributes to the increasing evidence that metagenomic NGS can effectively be used for a wide variety of diagnostic assays in virology, such as unbiased virus detection, resistance mutations, virulence markers, and epidemiology, as shown by the ability to detect SNPs in influenza virus.

These findings support the feasibility of moving this promising field forward to a role in the routine detection of pathogens by the use of mNGS. Further optimization should include the parallel evaluation of adult samples, the inclusion of additional annotated strain sequences to the database, and further elaboration of the classification algorithm and reporting for clinical diagnostics. The importance of both negative non-template control samples [41] and healthy control cases may support the critical discrimination of contaminants and viral ‘colonization’ from clinically relevant pathogens.

## CONCLUSIONS

Optimal sample preparation and bioinformatics analysis are essential for sensitive and specific mNGS based virus detection.

Using a high-throughput genome extraction method without viral enrichment, both RNA and DNA viruses could be detected with a sensitivity comparable to PCR.

Using mNGS, all potential pathogens can be detected in one single test, while simultaneously obtaining additional detailed information on detected viruses. Interpretation of clinical relevance is an important issue but essentially not different from the use of PCR based assays and supported by the available information on typing and relative quantities. These findings support the feasibility of a role of mNGS in the routine detection of pathogens.

## ACKNOWLEDGEMENTS

We thank our project partners Floyd Wittink, Wouter Suring (Hogeschool Leiden), Danny Duijsings (BaseClear) and Christiaan Henkel (Leiden University). We also like to thank thank Tom Vreeswijk, Lopje Höcker and Mario van Bussel (KML, LUMC) for help with the pre-sequencing experiments and Jeroen Laros (Human Genetics, LUMC) for help with the bioinformatics.

## AUTHOR CONTRIBUTIONS

SB, ALR, ECJC, ACMK, and JJCV participated in the study design. SB performed the pre-library preparation experiments. SB, NP, ECC, RHPV, PH, and HM carried out bioinformatic analyses. SB, ALR and ECC analyzed the data. SB and ALR wrote the first version of the manuscript. All authors contributed and revised the manuscript and approved the final manuscript.

## DATA ACCESS

The raw datasets of this study are not publicly made available given the confidential character of human sequences.

